# Neutrophils in patients with chronic coronary syndrome exhibit delayed spontaneous apoptosis and resistance to regulatory T cell-induced apoptosis – Brief Report

**DOI:** 10.64898/2026.01.22.701200

**Authors:** Maike Schneider, Camilla Skoglund, Rosanna WS Chung, Lena Jonasson

**Author notes:** Corresponding author: Lena Jonasson, Division of Diagnostics and Specialist Medicine, Department of Health, Medicine, and Caring Sciences, Linköping University, SE-58183 Linköping, Sweden.

## Abstract

**Background:** Persistent inflammation is linked to poor outcomes in patients with chronic coronary syndrome (CCS). The inflammatory state in atherosclerotic disease has been associated with activation of neutrophils as well as with regulatory T cell (T_reg_) deficiency. The role of T_regs_ in the regulation of neutrophil survival has been postulated recently. Here, we investigated neutrophil apoptosis along with the potential impact of T_regs_ on neutrophil apoptosis in patients with CCS, compared to healthy controls.

**Methods:** Twenty patients with CCS and 19 healthy controls were included. Neutrophil apoptosis was assessed after 5h culture with or without interleukin(IL-)10, TNF or LPS. Neutrophil phenotype was evaluated through flow cytometry analysis of surface receptors (CD66b and CXCR4) and *ex vivo* release of cytokines and granule proteins. The ability of T_regs_ to induce neutrophil apoptosis was examined in autologous neutrophil-T_reg_ co-cultures.

**Results:** Spontaneous neutrophil apoptosis was significantly delayed in patients compared to controls (10.3% vs. 19.2%, p=0.025). Also, neutrophils from patients overexpressed CD66b and CXCR4 and were more prone to release proinflammatory mediators. Notably, T_regs_ induced neutrophil apoptosis in healthy subjects, but not in patients, indicating a loss of T_reg_-mediated regulation in the latter. There was no evidence that IL-10 had any influence on neutrophil apoptosis. However, cell-to-cell contact was found essential for T_reg_-induced neutrophil apoptosis.

**Conclusions:** Patients with CCS display delayed neutrophil apoptosis and a pro-inflammatory neutrophil phenotype that is resistant to T_reg_-mediated apoptosis. Neutrophil dysfunction may contribute to persistent inflammation in patients with CCS, and as such constitute a novel therapeutic target.

## Introduction

The relationship between inflammation and atherosclerosis is well established, as shown in experimental, epidemiological and clinical studies ^1^. Despite optimal management of classical risk factors like hyperlipidemia and hypertension, systemic low-grade inflammation promotes recurrent cardiovascular events in patients with chronic coronary syndrome (CCS), a condition former known as stable coronary artery disease ^2^.

Growing evidence implicates neutrophils as pivotal mediators in atherosclerotic inflammation ^3^. Epidemiological studies consistently link elevated neutrophil counts to incident atherosclerotic cardiovascular disease and mortality ^4^. Increased neutrophil activation is also a well-known feature of the acute coronary syndrome (ACS) ^5–7^, while the occurrence of neutrophil activation in patients with CCS has been less studied. However, elevated neutrophil counts as well as neutrophil granule proteins, including myeloperoxidase (MPO), matrix metalloprotease (MMP)-8 and MMP-9, have been reported in patients with CCS, indicating a heightened neutrophil activation status ^8–11^. A few previous studies have also investigated whether neutrophils from patients with CCS are more prone to activation *ex vivo* compared to neutrophils from healthy controls, although with inconsistent results ^12–14^.

Neutrophils have a very short life, around 6 h, in circulating blood, They are inherently programmed to migrate back to the bone marrow via CXCR4 expression and die by constitutive apoptosis ^15^. Neutrophil apoptosis is considered to be a critical mechanism for resolving acute inflammation. A delay in the apoptotic process has been associated with increased neutrophil effector functions ^16,17^. Accordingly, delayed neutrophil apoptosis has been observed in a number of chronic inflammatory disorders ^18–20^. A few previous studies have also described delayed neutrophil apoptosis in patients with coronary artery disease, though with focus on ACS ^21,22^. Whether a delay in neutrophil apoptosis occurs in the stabilized phase after an ACS has not been clarified.

In patients with CCS, reduced numbers as well as impaired function of regulatory T cells (T_regs_) have been a consistent finding ^23–25^. T_regs_ are primarily recognized for their ability to modulate adaptive immunity. However, it has also been demonstrated that murine as well as human T_regs_ are able to promote neutrophil apoptosis and inhibit neutrophil activity ^26,27^. Interestingly, the role of dysregulated neutrophil-T_reg_ interaction in chronic inflammatory diseases, such as rheumatoid arthritis and systemic lupus erythematosus, has been proposed ^28,29^. Whether a dysregulated neutrophil-T_reg_ interaction takes place in patients with CCS has not previously been explored.

In the present study, we hypothesized that delayed neutrophil apoptosis is a feature of the persistent inflammatory state in patients with CCS. We examined neutrophil apoptosis, along with neutrophilś activation/aging status and their susceptibility to autologous T_regs_, in patients with CCS and healthy control subjects.

## Methods

The data that support the findings of this study are available from the corresponding author upon reasonable request. A detailed, expanded method sectionis is available in the Data Supplement.

### Study population

Twenty patients with CCS were recruited; all had angiographically verified coronary artery disease and a prior history of ACS. Patients were not eligible if they were more than 75 years of age, suffered from severe heart failure, immunological disorders, neoplastic disease, received treatment with immunosuppressive or anti-inflammatory agents (except low-dose aspirin), or had a history of acute or recent (<2 months) infection, major trauma/surgery or any revascularization procedure. For the control group (n = 19), volunteers with generally equal age and sex distribution were randomly selected from the Swedish Population Register. The study was conducted in accordance with the ethical guidelines of the Declaration of Helsinki, and the protocol was approved by the Ethical Review Board of Linkoping. Written informed consent was obtained from all study participants.

### Cell isolation and culture

Venous blood was collected in the morning in a fasting state. CD4^+^CD25^+^CD127^-/low^ T_regs_ were isolated from peripheral blood mononuclear cells (PBMCs) using an EasySep Human CD4+CD127lowCD25+ Regulatory T Cell Isolation Kit, and CD4+ T cells were isolated from PBMCs using a CD4+ T Cell Isolation Kit human. Neutrophils were isolated from whole blood using Polymorphprep.

For immediate staining, neutrophils were incubated with mouse anti-human CXCR4 APC and mouse anti-human CD66b PE antibodies for 15 min (Table S1). They were analysed within 1h using a Gallios Flow Cytometer.

Pure neutrophils, or co-cultures with neutrophils and T_regs_ or CD4^+^ T cells were seeded in 96-well plates at concentrations of 0.5 x10^6^ cells/ml for 2.5 h or 5 h, with or without interleukin (IL)-10, tumor necrosis factor (TNF) or lipopolysaccharide from *E. coli* (LPS).

### Neutrophil apoptosis and activation

Neutrophils were incubated for either 2.5 h or 5 h in different series of experiments. After 2.5 h, supernatants were stored at -80°C until further analysis. After 5 h, cells were stained with mouse anti-human CD66b PE, anti-human Annexin V BV42 and mouse anti-human CD4 BV510 antibodies (if the sample contained T_regs_ or CD4^+^ cells) in Annexin V Binding Buffer, followed by staining with SYTOX Red dead cell stain. Cells were analysed within 1 h using a Gallios Flow Cytometer to monitor apoptosis and activation status of neutrophils. Gating strategies are shown in Figure S1.

### Transwell experiments

Isolated neutrophils and T_regs_ were seeded in a 48-well cell culture plate with or without Millicell Cell Culture Inserts.

### Proteins in plasma and supernatants

Tumor necrosis factor (TNF), interferon(IFN)-γ, interleukin(IL)-6, IL-1β and IL-18 were measured in plasma via Olink Target 48, with an inter-assay CV of 7% and an intra-assay CV of 6%. Matrix metalloproteinase(MMP)-9, MMP-8, myeloperoxidase (MPO) in plasma were measured using a Luminex Multiplex Assay, with an intra-assay CV of 3.2%. MMP-9, MMP-8, MPO, TNF, IFN-γ, IL-6, IL-10, IL-1β and IL-18 were also measured in supernatants using a Luminex Multiplex Assay, with an inter-assay CV of 16.6% and an intra-assay CV of 8.1%. Measuements were performed by the Affinity Proteomics-Unit of SciLifeLab, Stockholm, Sweden.

### Statistics

All statistics were performed using the software IBM SPSS Statistics version 29.0.2.0. For comparisons between groups, a Mann-Whitney U test was performed. For comparison of paired data from the same individual, a Wilcoxon signed rank test was performed. For analysis of concentration-dependency, the Greenhouse-Geisser Test was performed.

## Results

### Cohort characteristics

All patients with CCS were clinically and metabolically stable with no symptoms of angina. There were no differences between patients and controls regarding age (median age 65 *vs* 64 years), sex distribution (females 50 % *vs* 53 %), body mass index or smoking status. Also, the protein levels in plasma were comparable between groups, except for higher levels of TNF and IL-10, as well as slightly but not significantly higher levels of IL-6, in the CCS group. Blood samples were collected from patients at a median of 11.5 months (interquartile range: 5.8-13 months) after an ACS. Cohort characteristics are presented in Table S2.

### Neutrophil apoptosis is delayed in patients with CCS

The apoptotic rate of neutrophils was assessed after 5 h culture in medium only (spontaneous apoptosis) or in the presence of either anti-inflammatory (IL-10) or pro-inflammatory (TNF, LPS) stimuli. As shown in Figure 1A, the spontaneous apoptosis was significantly reduced in CCS patients compared to healthy controls (10.3% vs. 19.2%, *p* = 0.025). A similar result was seen after 5 h treatment with IL-10 (10.1% vs. 19.8%, *p* = 0.020). Under inflammatory conditions (TNF or LPS), the differences in neutrophil apoptosis between patients and controls did not reach statistical significance.

**Figure 1:**
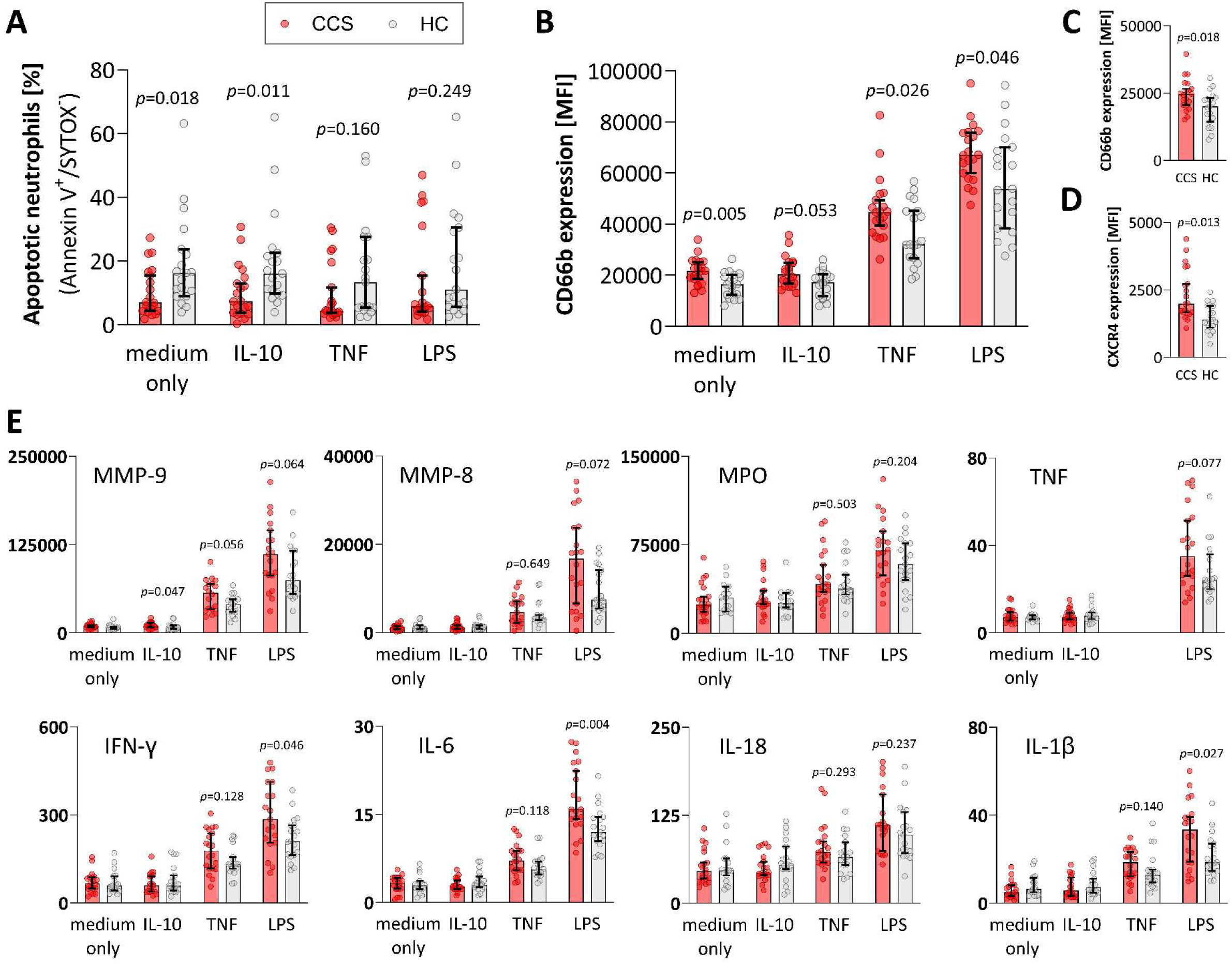
Neutrophil apoptosis, cell surface marker expression and neutrophil activation in patients with chronic coronary syndrome and healthy controls. Isolated neutrophils from 20 patients with chronic coronary syndrome (CCS) and 19 healthy controls (HC) were cultured in medium only or medium with interleukin (IL)-10 (20 ng/ml), tumor necrosis factor (TNF, 1 ng/ml) or lipopolysaccharide (LPS, 100 ng/ml). After 5 h, flow cytometry was used to assess neutrophil apoptosis; data are presented as the percentage of apoptotic (Annexin^+^/SYTOX^-^) neutrophils (**A**). Flow cytometry was also used to assess the expression of CD66b; data are presented as mean fluorescence intensity (MFI) values (**B**). Furthermore, the neutrophil expression of CD66b and CXCR4 was assessed by flow cytometry immediately after cell isolation in the same study cohort. Data are presented as MFI values of CD66b (**C**) and MFI values of CXCR4 (**D**). Additionally, protein concentrations were measured in the same study cohort after 2.5 h of incubation in medium with or without IL-10 (20 ng/ml), TNF (1 ng/ml) or LPS (100 ng/ml) (**E**). Concentrations of matrix metalloprotease (MMP)-9, MMP-8 and myeloperoxidase (MPO), TNF, interferon (IFN)γ, IL-6, IL-18 and IL-1β are presented as pg/ml. Significances between CCS and HC in normal medium and IL-10 were all *p* > 0.05, except for MMP-9 measurement in IL-10 treated sample. Significances are indicated between CCS and HC and all values are given as median ± 95% confidence interval.

### Neutrophils exhibit an aged and pro-inflammatory phenotype in patients with CCS

The surface expression of CD66b was used as a marker of neutrophil activation ^30^. After 5 h in medium only or in the presence of IL-10, TNF, or LPS the expression of CD66b was significantly higher in patients than in controls (Figure 1B).

The surface expression of CD66b and CXCR4, the latter representing neutrophil aging ^15^, was also assessed immediately after cell isolation. Neutrophils from CCS patients exhibited increased surface expression of CD66b as well as increased expression of CXCR4, compared to neutrophils from healthy subjects (Figure 1C-D).

To further investigate the neutrophil activation status, we measured the release of neutrophil granule proteins (MMP-9, MMP-8, MPO) and cytokines (TNF, IFN-γ, IL-6, IL-18, IL-1β) after a shorter period of incubation (2.5 h) with or without IL-10, TNF or LPS. There were no differences between patients and controls when neutrophils were cultured in medium only. However, upon LPS stimulation, the neutrophils from patients secreted higher levels of MMP-9, MMP-8, TNF, IFN-γ, IL-6 and IL-1β compared to controls. Also, the neutrophils from patients released higher levels of MMP-9 upon TNF treatment (Figure 1E).

### T_regs_ induce neutrophil apoptosis in healthy controls, but not in patients with CCS

We then proceeded to examine the effect of autologous T_regs_ on neutrophil apoptosis by co-culturing the cells at varying ratios (neutrophils:T_regs_ 1:0, 10:1, 4:1 and 1:1) for 5 h with or without IL-10, TNF or LPS. In the controls, T_regs_ induced neutrophil apoptosis in a concentration-dependent manner under all conditions, whereas no such effect was observed in the patients (Figure 2A-B). When T_reg_-induced neutrophil apoptosis was compared between patients and controls, the controls exhibited a significantly higher apoptotic rate at 5 h (with or without IL-10) at all ratios (Figures S2A-B). Autologous CD4^+^ T cells did not induce neutrophil apoptosis, neither in healthy controls nor in CCS patients (Figure S2A-D for between-groups analysis, Figure S3 for within-group analysis).

**Figure 2:**
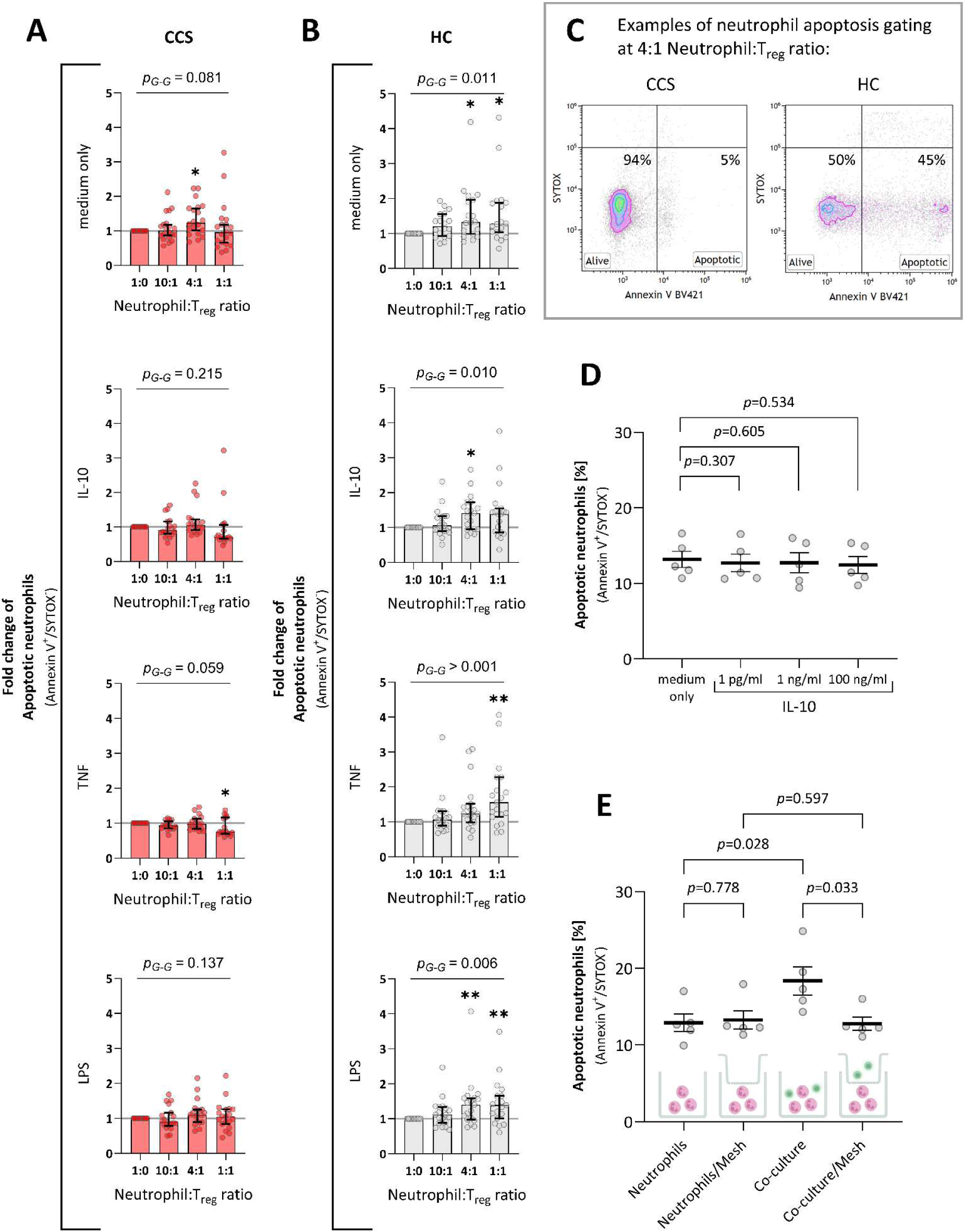
The efect of neutrophil-T_reg_ co-cultures on neutrophil apoptosis in patients with chronic coronary syndrome and healthy controls. Neutrophils and T_regs_ were isolated from 20 patients with chronic coronary syndrome (CCS) and 19 healthy controls (HC). The cells were co-cultured at diferent ratios (neutrophils: T_regs_ 1:0, 10:1, 4:1 and 1:1) in medium only or medium with interleukin (IL)-10 (20 ng/ml), tumor necrosis factor (TNF, 1 ng/ml) or lipopolysaccharide (LPS, 100 ng/ml). After 5 h of co-culture, neutrophil apoptosis (Annexin^+^/SYTOX^-^) was determined by flow cytometry. The percentages of apoptotic neutrophils are presented as fold change compared to pure neutrophil culture in patients with CCS (**A**) and HC (**B**). Significances are compared to 1:0 culture within CCS and HC groups, respectively, *p*-values are adjusted for multiple comparison via Dunn; CCS (**A**): \**p*=0.015 in medium only ratio 4:1, \**p*=0.036 in TNF ratio 1:1; HC (**B**): \**p*=0.014 in medium only ratio 4:1, \**p*=0.030 in medium only ratio 1:1, \**p*=0.025 in IL-10 ratio 4:1, \*\**p*=0.002 in TNF ratio 1:1, \*\**p*=0.009 in LPS ratio 4:1, \*\**p*=0.006 in LPS ratio 1:1; and significance for trend is given as *p*-value of a Greenhouse-Geisser test. All values are given as median ± 95% confidence interval. Representative examples for the gating of apoptotic neutrophils as Annexin V^+^/SYTOX^-^ cells are shown in **C**, the full gating strategy can be reviewed in Figure S1. Additionally, isolated neutrophils from healthy volunteers (n=5, < 50 years of age) were treated with diferent doses of IL-10 (1 pg/ml, 1ng/ml and 100ng/ml). Flow cytometry was used to assess neutrophil apoptosis after 5 h; data are presented as the percentage of apoptotic (Annexin^+^/SYTOX^-^) neutrophils (**D**). In another series of experiments, using cells from the same volunteers described in (**D**), neutrophils alone or neutrophil-T_reg_ co-cultures at ratio 4:1 were seeded +/- mesh (as graphically indicated) in medium only for 5 h, followed by flow cytometry to assess neutrophil apoptosis (**E**). *p*-values were adjusted for multiple comparison via Tukey. All values are given as mean ± SEM.

### T_reg_ survival and IL-10 secretion in neutrophil-T_reg_ co-cultures do not differ between patients with CCS and healthy controls

To determine whether differences in T_reg_ survival contributed to the discrepancy in T_reg_-induced neutrophil apoptosis between groups, we assessed the T_reg_ viability under various conditions. Neither T_reg_ nor CD4^+^ T cell survival differed between patients and controls under any conditions (Figure S4A-B).

To gain an insight into the functionality of T_regs_, IL-10 was measured in co-culture supernatants of patients and controls. In controls, the concentrations of IL-10 appeared to increase at ratios 4:1 and 1:1 (Figure S4C-D). However, the absolute levels were low (0.2 - 1.1 pg/ml), and no significant differences were observed between patients and controls.

### T_reg_-induced neutrophil apoptosis is not dependent on IL-10, but rather on cell-to-cell contact

Next, we investigated two possible mechanisms of T_reg_-induced neutrophil apoptosis. To determine whether IL-10 was a potential mediator of T_reg_-induced neutrophil apoptosis, neutrophils from healthy volunteers were treated with increasing concentrations of IL-10 (1 pg/ml, 1 ng/ml, 100 ng/ml) for 5 h, mimicking concentrations in co-culture supernatants (mentioned above) as well as concentrations used in previous *in vitro* studies ^31,32^. None of the IL-10 concentrations showed any effect on the apoptotic rate (Figure 2D). To further assess whether T_regs_ used cell-to-cell contact to regulate neutrophil apoptosis, transwell experiments were performed to physically separate T_regs_ and neutrophils via a membrane (mesh) at ratio 4:1. As shown in Figure 2E, the separation completely abrogated T_reg_-induced neutrophil apoptosis.

## Discussion

The main finding of the present study is that spontaneous neutrophil apoptosis, a key process in the resolution of inflammation, was significantly delayed in patients with CCS compared to healthy controls. Concomitantly, the neutrophils in patients exhibited an aged and proinflammatory phenotype as well as a resistance to T_reg_-induced neutrophil apoptosis.

Delayed neutrophil apoptosis has been described in several chronic inflammatory diseases ^18–20^. Only a few earlier studies have investigated neutrophil death in coronary artery disease, with emphasis on ACS, a condition characterized by acute inflammation. In 2004, Garlich *et al.* ^21^ demonstrated that neutrophil death, including both apoptosis and necrosis, was delayed in patients with ACS (n=30), compared to patients with stable angina (n=15) and controls (n=15) after 24 h culture. Some years later, Biasucci *et al.* ^22^ measured spontaneous neutrophil apoptosis in patients with ACS (n=30), patients with stable angina (n=13) and controls (n=34). They found delayed neutrophil apoptosis in ACS patients after 4 h culture (10.1%), whereas the apoptotic rate in stable angina patients (28.5%) did not differ from that in controls (41.8%). In the present study, we detected a delay in neutrophil apoptosis already after 5 h in patients with CCS compared to controls. Comparisons with earlier study groups should however be drawn with caution, e.g. the apoptotic rates reported by Biasucci *et al.* ^22^ after 4 h culture were remarkably higher than what we found, which might indicate methodological differences. Also, we included patients who had survived ACS, while Biasucci *et al.* ^22^ included stable angina patients without prior ACS. Despite being asymptomatic and without overt heart failure, it cannot be excluded that our patient cohort had a more severe, complicated coronary artery disease, hence a more proinflammatory neutrophil phenotype.

It is well established that patients with ACS exhibit a more active neutrophil phenotype, either within culprit coronary lesions or circulating in peripheral blood ^3^. Whether the active neutrophil phenotype is maintained in those stabilized after a prior ACS has been less clear. Results from *ex vivo* functional assays are sparse; some have reported that circulating neutrophils in patients with CCS are primed, i.e. more prone to stimulation *ex vivo*, as compared to neutrophils in healthy controls ^12,13^, while others have not been able to detect any differences ^14,33^. In the present study, delayed spontaneous neutrophil apoptosis in patients with CCS was clearly accompanied by increased expression of CD66b, a sensitive marker of neutrophil activation in human samples ^30^. Also, the finding that patientśneutrophils were more prone to secrete cytokines and granule proteins upon stimulation *ex vivo* supported their state of heightened activity. Furthermore, the increased expression of CXCR4 indicate an accumulation of aged neutrophils in the circulation of patients with CCS. Aged neutrophils have been shown to exhibit a distinct, highly reactive phenotype, both in experimental models and in chronic inflammatory disease ^34,35^.

Statins are known to exert pleiotropic anti-inflammatory effects, one of which is the suppression of neutrophil activation ^36,37^. Furthermore, two single-center randomized trials have shown that statins promote neutrophil apoptosis in acute inflammatory conditions ^38,39^. Despite extensive treatment with statins, CCS patients in our study exhibited an increase in proinflammatory neutrophils with delayed apoptosis. These findings indicate that statin-mediated immunomodulatory effects on innate immunity, while robust *in vitro*, may be less effective or relevant in chronic inflammation.

T_regs_ are well known for their role in controling the adaptive immune response ^29^. Emerging evidence indicates that T_regs_ also have a regulatory role in the innate immune response ^29^. Of note, a deficit in circulating T_regs_ is a characteristic finding in patients with both acute and chronic manifestations of coronary artery disease ^23–25^, as well as in patients with other chronic inflammatory diseases ^40–42^. Earlier experimental studies, mainly performed on murine cells, have shown that LPS-activated T_regs_ are able to inhibit neutrophil function and promote neutrophil apoptosis ^26,27^. In the present study, T_regs_ promoted neutrophil apoptosis both with and without LPS in healthy controls, while this phenomenon did not occur in patients with CCS. The reason for resistance to T_reg_-induced neutrophil apoptosis may be multifactorial, depending on anti-apoptotic features of pro-inflammatory neutrophils ^43^ but also on functional defects of T_regs_. As previously described by our group, the capacity of T_regs_ to regulate proliferation and secretion of IFN-γ and IL-10 in autologous T responder cells is impaired in patients with CCS compared to healthy controls ^25^. Several studies have also shown that a chronic inflammatory environment may lead to impaired T_reg_ function depending on molecular mechanisms, such as perturbed *Foxp3* expression and epigenetic modifications ^44^.

IL-10 has been described as a mediator of anti-inflammatory effects exerted by T_regs_ ^45,46^. In the present study, we used a wide range of concentrations of IL-10 in both neutrophil cultures and neutrophil-T_reg_ co-cultures, but were not able to demonstrate any effect on neutrophil apoptosis after 5 h. This is in agreement with earlier studies reporting that IL-10 does not affect spontaneous neutrophil apoptosis, neither after short (6 h) nor long (20 h) incubation ^47,48^. However, these earlier studies have also reported that, under *in vitro* conditions mimicking severe sepsis, IL-10 is capable of counteracting LPS-induced delay of neutrophil apoptosis ^31,47,48^. Moreover, LPS-stimulated T_regs_ have been shown to induce IL-10 production in apoptotic neutrophils after 5 h co-culture, while LPS alone or unstimulated T_regs_ do not ^49^. In the present study, we detected very low concentrations of IL-10 in supernatants after 5 h co-culture, both with and without LPS, speaking against a significant role of IL-10 in this experimental model for T_reg_-induced neutrophil apoptosis. On the other hand, our findings clearly show that T_reg_-induced neutrophil apoptosis was dependent on cell-to-cell contact. The results are in agreement with previous studies reporting that human T_regs_ exert suppressive effects through cell-to-cell contact ^49,50^.

Several limitations of our study should be acknowledged. The neutrophil-T_reg_ co-cultures were performed at cell ratios that do not reflect physiological conditions in peripheral blood where the neutrophil:T_regs_ ratio is normally 100:1. However, the *in vitro* approach using ratios from 1:1 to 10:1 is in line with previous mechanistic studies ^26,27,51^. Another major limitation is that we did not measure the phenotype of isolated T_regs_, which limits the understanding of why neutrophil-T_reg_ interaction was impaired in patients compared to controls. Lastly, the sample size was small and our findings should be considered preliminary until validated in a larger cohort.

Nevertheless, a strength of the study is the *ex vivo* investigation of peripheral neutrophils in human disease, addressing a critical gap in translational cardiovascular research. While numerous epidemiological studies have shown that high neutrophil counts is a causal risk factor for cardiovascular disease ^4^, mechanistic insights have been mainly derived from animal models ^3^.

To conclude, delayed spontaneous neutrophil apoptosis, along with a more active neutrophil phenotype showing resistance to T_reg_-induced apoptosis, indicate that neutrophils may be a major contributor to the non-resolved inflammatory state in patients with CCS. Of note, neutrophil dysfunction was observed in clinically stable patients who received optimal medical therapy according to current guidelines. Although the findings need to be confirmed in larger samples, and also explored in further mechanistic studies, they allow us to speculate about neutrophil apoptosis as a future target of therapy in patients with CCS.

## Acknowledgements

We acknowledge the Core Facility at the Faculty of Medicine and Health Sciences, Linköping University for providing assistance in flow cytometry and the Affinity Proteomics Unit at SciLifeLab in Stockholm for supporting with protein quantification.

## References

1. Libby P, Hansson GK. From Focal Lipid Storage to Systemic Inflammation: JACC Review Topic of the Week. Journal of the American College of Cardiology. 2019;74:1594–1607. doi: 10.1016/j.jacc.2019.07.061

2. Libby P, Soehnlein O. Inflammation in atherosclerosis: Lessons and therapeutic implications. Immunity. 2025;58:2383–2401. doi: 10.1016/j.immuni.2025.09.012

3. Döring Y, Drechsler M, Soehnlein O, Weber C. Neutrophils in Atherosclerosis. Arteriosclerosis, Thrombosis, and Vascular Biology. 2015;35:288–295. doi: doi:10.1161/ATVBAHA.114.303564

4. Luo J, Thomassen JQ, Nordestgaard BG, Tybjærg-Hansen A, Frikke-Schmidt R. Neutrophil counts and cardiovascular disease. European Heart Journal. 2023;44:4953–4964. doi: 10.1093/eurheartj/ehad649

5. Bonaventura A, Montecucco F, Dallegri F, Carbone F, Lüscher TF, Camici GG, Liberale L. Novel findings in neutrophil biology and their impact on cardiovascular disease. Cardiovasc Res. 2019;115:1266–1285. doi: 10.1093/cvr/cvz084

6. Talmor N, Pillinger MH, Xia Y, Leonard A, Curovic F, Shah B. Neutrophil Activation and Adhesiveness in Coronary Artery Disease: Results From the COLCHICINE&#x2010;PCI Biomarker Substudy. Journal of the American Heart Association. 2024;13:e036701. doi: doi:10.1161/JAHA.124.036701

7. Maréchal P, Tridetti J, Nguyen ML, Wéra O, Jiang Z, Gustin M, Donneau AF, Oury C, Lancellotti P. Neutrophil Phenotypes in Coronary Artery Disease. J Clin Med. 2020;9. doi: 10.3390/jcm9051602

8. Momiyama Y, Ohmori R, Tanaka N, Kato R, Taniguchi H, Adachi T, Nakamura H, Ohsuzu F. High plasma levels of matrix metalloproteinase-8 in patients with unstable angina. Atherosclerosis. 2010;209:206–210. doi: 10.1016/j.atherosclerosis.2009.07.037

9. Caselli C, Di Giorgi N, Ragusa R, Lorenzoni V, Smit J, El Mahdiui M, Buechel RR, Teresinska A, Pizzi MN, Roque A, et al. Association of MMP9 with adverse features of plaque progression and residual inflammatory risk in patients with chronic coronary syndrome (CCS). Vascul Pharmacol. 2022;146:107098. doi: 10.1016/j.vph.2022.107098

10. Nicholls SJ, Hazen SL. Myeloperoxidase and Cardiovascular Disease. Arteriosclerosis, Thrombosis, and Vascular Biology. 2005;25:1102–1111. doi: doi:10.1161/01.ATV.0000163262.83456.6d

11. Avanzas P, Arroyo-Espliguero R, Cosín-Sales J, Quiles J, Zouridakis E, Kaski JC. Multiple complex stenoses, high neutrophil count and C-reactive protein levels in patients with chronic stable angina. Atherosclerosis. 2004;175:151–157. doi: 10.1016/j.atherosclerosis.2004.03.013

12. Paulsson J, Dadfar E, Held C, Jacobson SH, Lundahl J. Activation of peripheral and in vivo transmigrated neutrophils in patients with stable coronary artery disease. Atherosclerosis. 2007;192:328–334. doi: 10.1016/j.atherosclerosis.2006.08.003

13. Jönsson S, Lundberg A, Kälvegren H, Bergström I, Szymanowski A, Jonasson L. Increased Levels of Leukocyte-Derived MMP-9 in Patients with Stable Angina Pectoris. PLOS ONE. 2011;6:e19340. doi: 10.1371/journal.pone.0019340

14. Särndahl E, Bergström I, Brodin VP, Nijm J, Lundqvist Setterud H, Jonasson L. Neutrophil activation status in stable coronary artery disease. PLoS One. 2007;2:e1056. doi: 10.1371/journal.pone.0001056

15. Martin C, Burdon PC, Bridger G, Gutierrez-Ramos JC, Williams TJ, Rankin SM. Chemokines acting via CXCR2 and CXCR4 control the release of neutrophils from the bone marrow and their return following senescence. Immunity. 2003;19:583–593. doi: 10.1016/s1074-7613(03)00263-2

16. Brostjan C, Oehler R. The role of neutrophil death in chronic inflammation and cancer. Cell Death Discovery. 2020;6:26. doi: 10.1038/s41420-020-0255-6

17. Ocaña MG, Asensi V, Montes AH, Meana A, Celada A, Valle-Garay E. Autoregulation mechanism of human neutrophil apoptosis during bacterial infection. Mol Immunol. 2008;45:2087–2096. doi: 10.1016/j.molimm.2007.10.013

18. Midgley A, McLaren Z, Moots RJ, Edwards SW, Beresford MW. The role of neutrophil apoptosis in juvenile-onset systemic lupus erythematosus. Arthritis Rheum. 2009;60:2390–2401. doi: 10.1002/art.24634

19. Weinmann P, Moura RA, Caetano-Lopes JR, Pereira PA, Canhão H, Queiroz MV, Fonseca JE. Delayed neutrophil apoptosis in very early rheumatoid arthritis patients is abrogated by methotrexate therapy. Clin Exp Rheumatol. 2007;25:885–887.

20. Brannigan AE, O’Connell PR, Hurley H, O’Neill A, Brady HR, Fitzpatrick JM, Watson RW. Neutrophil apoptosis is delayed in patients with inflammatory bowel disease. Shock. 2000;13:361–366. doi: 10.1097/00024382-200005000-00003

21. Garlichs CD, Eskafi S, Cicha I, Schmeisser A, Walzog B, Raaz D, Stumpf C, Yilmaz A, Bremer J, Ludwig J, et al. Delay of neutrophil apoptosis in acute coronary syndromes. J Leukoc Biol. 2004;75:828–835. doi: 10.1189/jlb.0703358

22. Biasucci LM, Liuzzo G, Giubilato S, Della Bona R, Leo M, Pinnelli M, Severino A, Gabriele M, Brugaletta S, Piro M, et al. Delayed neutrophil apoptosis in patients with unstable angina: relation to C-reactive protein and recurrence of instability. Eur Heart J. 2009;30:2220–2225. doi: 10.1093/eurheartj/ehp248

23. Mor A, Luboshits G, Planer D, Keren G, George J. Altered status of CD4(+)CD25(+) regulatory T cells in patients with acute coronary syndromes. Eur Heart J. 2006;27:2530–2537. doi: 10.1093/eurheartj/ehl222

24. George J, Schwartzenberg S, Medvedovsky D, Jonas M, Charach G, Afek A, Shamiss A. Regulatory T cells and IL-10 levels are reduced in patients with vulnerable coronary plaques. Atherosclerosis. 2012;222:519–523. doi: 10.1016/j.atherosclerosis.2012.03.016

25. Hasib L, Lundberg AK, Zachrisson H, Ernerudh J, Jonasson L. Functional and homeostatic defects of regulatory T cells in patients with coronary artery disease. J Intern Med. 2016;279:63–77. doi: 10.1111/joim.12398

26. Lewkowicz P, Lewkowicz N, Sasiak A, Tchórzewski H. Lipopolysaccharide-activated CD4+CD25+ T regulatory cells inhibit neutrophil function and promote their apoptosis and death. J Immunol. 2006;177:7155–7163. doi: 10.4049/jimmunol.177.10.7155

27. Okeke EB, Mou Z, Onyilagha N, Jia P, Gounni AS, Uzonna JE. Deficiency of Phosphatidylinositol 3-Kinase δ Signaling Leads to Diminished Numbers of Regulatory T Cells and Increased Neutrophil Activity Resulting in Mortality Due to Endotoxic Shock. J Immunol. 2017;199:1086–1095. doi: 10.4049/jimmunol.1600954

28. Okeke EB, Uzonna JE. The Pivotal Role of Regulatory T Cells in the Regulation of Innate Immune Cells. Front Immunol. 2019;10:680. doi: 10.3389/fimmu.2019.00680

29. Ou Q, Power R, Griffin MD. Revisiting regulatory T cells as modulators of innate immune response and inflammatory diseases. Front Immunol. 2023;14:1287465. doi: 10.3389/fimmu.2023.1287465

30. Ledderose C, Hashiguchi N, Valsami EA, Rusu C, Junger WG. Optimized flow cytometry assays to monitor neutrophil activation in human and mouse whole blood samples. J Immunol Methods. 2023;512:113403. doi: 10.1016/j.jim.2022.113403

31. Cox G. IL-10 enhances resolution of pulmonary inflammation in vivo by promoting apoptosis of neutrophils. Am J Physiol. 1996;271:L566–571. doi: 10.1152/ajplung.1996.271.4.L566

32. Laichalk LL, Danforth JM, Standiford TJ. Interleukin-10 inhibits neutrophil phagocytic and bactericidal activity. FEMS Immunol Med Microbiol. 1996;15:181–187. doi: 10.1111/j.1574-695X.1996.tb00084.x

33. Mazzone A, De Servi S, Ricevuti G, Mazzucchelli I, Fossati G, Pasotti D, Bramucci E, Angoli L, Marsico F, Specchia G, et al. Increased expression of neutrophil and monocyte adhesion molecules in unstable coronary artery disease. Circulation. 1993;88:358–363. doi: 10.1161/01.cir.88.2.358

34. Uhl B, Vadlau Y, Zuchtriegel G, Nekolla K, Sharaf K, Gaertner F, Massberg S, Krombach F, Reichel CA. Aged neutrophils contribute to the first line of defense in the acute inflammatory response. Blood. 2016;128:2327–2337. doi: 10.1182/blood-2016-05-718999

35. Chen J, Bai Y, Xue K, Li Z, Zhu Z, Li Q, Yu C, Li B, Shen S, Qiao P, et al. CREB1-driven CXCR4hi neutrophils promote skin inflammation in mouse models and human patients. Nature Communications. 2023;14:5894. doi: 10.1038/s41467-023-41484-3

36. Maher BM, Dhonnchu TN, Burke JP, Soo A, Wood AE, Watson RW. Statins alter neutrophil migration by modulating cellular Rho activity--a potential mechanism for statins-mediated pleotropic effects? J Leukoc Biol. 2009;85:186–193. doi: 10.1189/jlb.0608382

37. Antoniellis Silveira AA, Dominical VM, Morelli Vital D, Alves Ferreira W, Trindade Maranhão Costa F, Werneck CC, Ferreira Costa F, Conran N. Attenuation of TNF-induced neutrophil adhesion by simvastatin is associated with the inhibition of Rho-GTPase activity, p50 activity and morphological changes. International Immunopharmacology. 2018;58:160–165. doi: 10.1016/j.intimp.2018.03.025

38. Chello M, Anselmi A, Spadaccio C, Patti G, Goffredo C, Di Sciascio G, Covino E. Simvastatin increases neutrophil apoptosis and reduces inflammatory reaction after coronary surgery. Ann Thorac Surg. 2007;83:1374–1380. doi: 10.1016/j.athoracsur.2006.10.065

39. Mandal P, Chalmers JD, Graham C, Harley C, Sidhu MK, Doherty C, Govan JW, Sethi T, Davidson DJ, Rossi AG, et al. Atorvastatin as a stable treatment in bronchiectasis: a randomised controlled trial. Lancet Respir Med. 2014;2:455–463. doi: 10.1016/s2213-2600(14)70050-5

40. Li W, Deng C, Yang H, Wang G. The Regulatory T Cell in Active Systemic Lupus Erythematosus Patients: A Systemic Review and Meta-Analysis. Front Immunol. 2019;10:159. doi: 10.3389/fimmu.2019.00159

41. Jiang Q, Yang G, Liu Q, Wang S, Cui D. Function and Role of Regulatory T Cells in Rheumatoid Arthritis. Front Immunol. 2021;12:626193. doi: 10.3389/fimmu.2021.626193

42. Jalalvand M, Enayati S, Akhtari M, Madreseh E, Jamshidi A, Farhadi E, Mahmoudi M, Amirzargar A. Blood regulatory T cells in inflammatory bowel disease, a systematic review, and meta-analysis. Int Immunopharmacol. 2023;117:109824. doi: 10.1016/j.intimp.2023.109824

43. Noseykina EM, Schepetkin IA, Atochin DN. Molecular Mechanisms for Regulation of Neutrophil Apoptosis under Normal and Pathological Conditions. J Evol Biochem Physiol. 2021;57:429–450. doi: 10.1134/s0022093021030017

44. Dominguez-Villar M, Hafler DA. Regulatory T cells in autoimmune disease. Nature Immunology. 2018;19:665–673. doi: 10.1038/s41590-018-0120-4

45. Chaudhry A, Samstein RM, Treuting P, Liang Y, Pils MC, Heinrich JM, Jack RS, Wunderlich FT, Brüning JC, Müller W, et al. Interleukin-10 signaling in regulatory T cells is required for suppression of Th17 cell-mediated inflammation. Immunity. 2011;34:566–578. doi: 10.1016/j.immuni.2011.03.018

46. Laidlaw BJ, Cui W, Amezquita RA, Gray SM, Guan T, Lu Y, Kobayashi Y, Flavell RA, Kleinstein SH, Craft J, et al. Production of IL-10 by CD4+ regulatory T cells during the resolution of infection promotes the maturation of memory CD8+ T cells. Nature Immunology. 2015;16:871–879. doi: 10.1038/ni.3224

47. Keel M, Ungethüm U, Steckholzer U, Niederer E, Hartung T, Trentz O, Ertel W. Interleukin-10 counterregulates proinflammatory cytokine-induced inhibition of neutrophil apoptosis during severe sepsis. Blood. 1997;90:3356–3363.

48. Ward C, Murray J, Clugston A, Dransfield I, Haslett C, Rossi AG. Interleukin-10 inhibits lipopolysaccharide-induced survival and extracellular signal-regulated kinase activation in human neutrophils. Eur J Immunol. 2005;35:2728–2737. doi: 10.1002/eji.200425561

49. Lewkowicz N, Mycko MP, Przygodzka P, Ćwiklińska H, Cichalewska M, Matysiak M, Selmaj K, Lewkowicz P. Induction of human IL-10-producing neutrophils by LPS-stimulated Treg cells and IL-10. Mucosal Immunol. 2016;9:364–378. doi: 10.1038/mi.2015.66

50. Thornton AM, Shevach EM. CD4+CD25+ immunoregulatory T cells suppress polyclonal T cell activation in vitro by inhibiting interleukin 2 production. J Exp Med. 1998;188:287–296. doi: 10.1084/jem.188.2.287

51. Lewkowicz N, Klink M, Mycko MP, Lewkowicz P. Neutrophil--CD4+CD25+ T regulatory cell interactions: a possible new mechanism of infectious tolerance. Immunobiology. 2013;218:455–464. doi: 10.1016/j.imbio.2012.05.029

