## Supplementary material for "Neutrophils in patients with chronic coronary syndrome exhibit delayed spontaneous apoptosis and resistance to regulatory T cell-induced apoptosis – Brief Report": ATVB-2026-324652D_Schneider_SupplementalMaterial.docx

**Supplemental Material**

**Expanded method section**

Study population

Twenty patients with CCS were recruited from the Cardiology Outpatient Clinic at the University Hospital in Linköping, Sweden. All patients had angiographically verified CAD and a prior history of ACS. Patients were not eligible if they were more than 75 years of age, suffered from severe heart failure (defined as symptomatic or advanced heart failure with NYHA classifications III-IV), immunological disorders, neoplastic disease, received treatment with immunosuppressive or anti-inflammatory agents (except low-dose aspirin), or had a history of acute or recent (<2 months) infection, major trauma/surgery or any revascularization procedure. For the control group (n = 19), volunteers from Linköping with generally equal age and sex distribution were randomly selected from the Swedish Population Register and invited to participate in the study. Individuals who accepted the invitation were included as controls if they were anamnestically healthy and had normal routine laboratory tests. Use of lipid-lowering or antihypertensive drugs for primary prevention was allowed in the control group. The study was conducted in accordance with the ethical guidelines of the Declaration of Helsinki, and the research protocol was approved by the Ethical Review Board of Linkoping. Written informed consent was obtained from all study participants.

Cell isolation

Venous blood was always collected in the morning (between 8:00 and 9:30 AM ) in a fasting state. Peripheral blood mononuclear cells (PBMCs) were isolated from 40 ml whole blood in 4 10-ml sodium heparin-tubes (BD biosciences) diluted in PBS (1:1 [blood:PBS]) using Ficoll-Paque Density Gradient Medium (ThermoFisher Scientific. CD4^+^CD25^+^CD127^-/low^ T_regs_ were isolated from PBMCs using an EasySep Human CD4+CD127lowCD25+ Regulatory T Cell Isolation Kit (Stemcell Technologies), according to the manufacturer’s instructions. CD4+ T cells were isolated from PBMCs using a CD4+ T Cell Isolation Kit human (Miltenyi Biotec) according to the manufacturer’s instructions, resuspended in 500 µl RPMI Medium 1640 1x (Gibco) and kept on ice until seeding. Neutrophils were isolated from 20 ml EDTA whole blood using Polymorphprep (Fisher Scientific). Whole blood was carefully layered onto Polymorphprep (ratio 1:1) and centrifuged at 480xg for 40 min at room temperature (RT) with low acceleration and no deceleration. Thereafter, neutrophils were collected from the neutrophil phase with a transfer pipette (Sarstedt) and washed with 7.5 ml 0.45% NaCl (Sigma-Aldrich) in deionised H_2_O and 20 ml PBS, followed by centrifugation at 400xg for 10 min at RT. Erythrocytes were depleted using hypotonic lysis. Briefly, neutrophils were kept on ice and resuspended in 4.5 ml ice-cold water. After 35 seconds, the lysis was stopped by adding 1.5 ml of ice-cold 3.4% of NaCl in PBS and 5 ml of ice-cold KRG without Ca^2+^. Afterwards, neutrophils were centrifuged at 400xg for 5 min at 4 °C, washed in 20 ml PBS, centrifuged again at 400xg for 5 min at 4 °C, resuspended in 500 µl RPMI medium and kept on ice until seeding or immediate staining for CXCR4 and CD66b.

For immediate staining, 200,000 neutrophils were washed in 1 ml PBS + 0.05% fetal bovine serum (FBS, Gibco) centrifuged at 500xg for 5 min at RT and stained with mouse anti-human CXCR4 APC (12G5, BD Biosciences) and mouse anti-human CD66b PE (G10F5, BD Biosciences) antibodies (for concentrations see Table S1) in 60 µl PBS + 0.05% FBS. After 15 min of incubation at RT, cells were washed with PBS + 0.05% FBS, centrifuged at 500xg for 5 min at RT and resuspended in 400 µl PBS + 0.05% FBS. The neutrophils were analysed using a Gallios Flow Cytometer within 1h to monitor aging and activation status of neutrophils. For all analysis of flow cytometry data, the software Kaluza Analysis version 2.1.00002.20011 was used. For neutrophils, the purity determined via flow cytometry was 88.2% ± 4.4%. For T_regs_ and CD4^+^ T cells, the purity was 92.1% ± 6.2% and 97.0% ± 1.9%, respectively (values are presented as mean ± SD). There were no differences in purity of any isolated cell types between patients with CCS and healthy controls.

Cell culture

All cells were seeded in RPMI medium supplemented with 10% FBS and 2% Penicillin Streptomycin solution (Gibco). For some experiments lipopolysaccharides from *E. coli* (LPS, 026:B6, Sigma-Aldrich, 100 ng/ml), tumor necrosis factor (TNF/ Sigma-Aldrich, 1 ng/ml) or interleukin (IL)-10 (Sigma-Aldrich, 20 ng/ml or indicated concentration) were added to the medium. All cells were seeded in round-bottom 96-well plates (Sarstedt) at concentrations of 0.5 x10^6^ cells/ml. Pure neutrophils were seeded in 200 µl medium (0:1), while for co-cultures 20 µl or 50 µl of T_regs_ or CD4^+^ T cells (the latter used as control) were added to achieve a total of 220 µl and 250 µl of volume, resulting in the ratio 1:10 or 1:4, respectively. For the 1:1 cultures, 100 µl neutrophils and 100 µl T_regs_ or CD4^+^ T cells were seeded for a total of 200 µl per sample. All samples were incubated for 2.5 h or 5 h at standard conditions (37 °C and 5% CO_2_).

Neutrophil apoptosis and activation

Neutrophils were incubated for either 2.5 h or 5 h in different series of experiments. After 2.5 h of incubation, cells were centrifuged at 500xg for 5 min at RT and 150 µl supernatant per sample was stored at -80°C until further analysis. After 5 h of incubation, cells were centrifuged at 500xg for 5 min at RT, and thereafter washed in 600 µl PBS + 0.05% FBS, centrifuged at 500xg for 5 min at RT and stained with mouse anti-human CD66b PE (G10F5, BD biosciences), anti-human Annexin V BV421 (BD Biosciences), and mouse anti-human CD4 BV510 (SK3, BD Biosciences) antibodies (if the sample contained T_regs_ or CD4^+^ cells) (for concentrations see Supplementary table ST1) in 40 µl 1x Annexin V Binding Buffer (BD biosciences). After 15 min of incubation at RT, cells were washed with 560 µl Annexin V Binding Buffer, centrifuged at 500xg for 5 min at RT and resuspended in 200 µl Annexin V Binding Buffer. Cells were stained with SYTOX Red dead cell stain (25 nM final concentration, Thermofisher) and analysed using a Gallios Flow Cytometer within 1 h to monitor apoptosis and activation status of neutrophils. Gating strategies are shown in Figure S1.

Transwell experiments

Neutrophils and T_regs_ were isolated as described above and seeded in a 48-well cell culture plate (Falcon) with a cell concentration of 0.5 x10^6^ cells/ml. Four hundred µl of neutrophils were co-cultured with 100 µl of T_regs_ +/- Millicell Cell Culture Inserts, 0.4µm PET (Sigma Aldrich).

Measurement of proteins in plasma and supernatants

Blood in a 6 ml EDTA-tube (K2E, BD biosciences) was centrifuged, within 30 min of collection, at 1500xg for 10 min at RT. Plasma was aliquoted and frozen at -80 °C. TNF, interferon(IFN)γ, IL-6, IL-1β and IL-18 were measured via Olink Target 48, with an inter-assay CV of 7% and an intra-assay CV of 6%. MMP-8, MMP-9, MPO in plasma were measured using a customized Luminex Multiplex Assay (BD biosciences), with an intra-assay CV of 3.2%. CRP was measured at the Department of Clinical Chemistry at the Linköping University hospital using an immunoturbidimetric assay with a Roche Cobas C502 analyzer (Roche Diagnostics, Scandinavia AB), with a CV of 2.2%. MMP-8, MMP-9, MPO, TNF, IFN-γ, IL-6, IL-10, IL-1β and IL-18 in supernatants were measured using a customized Luminex Multiplex Assay (BD biosciences), with an inter-assay CV of 16.6% and an intra-assay CV of 8.1%. Measurements were performed by the Affinity Proteomics-Unit of SciLifeLab, Stockholm, Sweden.

Statistics

All statistics were performed using the software IBM SPSS Statistics version 29.0.2.0 or GraphPad Prism version 10.0.2. For comparisons between groups, a Mann-Whitney U test was performed. For comparison of paired data from the same individual, a test was performed. For analysis of concentration-dependency a test for repeated measurements, a Greenhouse-Geisser Test, was performed together with a Friedman test, followed by Dunn’s multiple comparisons test (for the percentage of neutrophil apoptosis in co-culture samples) or a Mixed-effects analysis, followed by Dunnett’s multiple comparisons test (IL-10 measurements). To compare different samples from the mechanistic experiments (n=5), a one-way ANOVA followed by Tukey’s multiple comparison test was performed.

**Table S1:** Antibody information

| **Staining** | **Antibody/stain** | **Fluoro-phore** | **Clone** | **Final concen-tration [µg/ml]** | **Cat nr.** | **Company** |
| --- | --- | --- | --- | --- | --- | --- |
| CXCR4  expression | mouse anti-human CXCR4 | APC | 12G5 | 5 | 555976 | BD Biosciences |
| CD66b expression | mouse anti-human CD66b | PE | G10F5 | 1.25 | 561650 | BD Biosciences |
| Apoptosis | anti-human Annexin V | BV421 | / | 2.6 | 563973 | BD Biosciences |
| Co-culture staining | mouse anti-human CD4 | BV510 | SK3 | 1.25 | 562970 | BD Biosciences |
| Purity check | mouse anti-human CD66b | PE | G10F5 | 2.5 | 561650 | BD biosciences |
| Purity check | mouse anti-human Siglec-8 | PE-Cy7 | 7C9 | 10 | 347112 | Biolegend |
| Purity check | mouse anti-human CD25 | PE | 2A3 | 0.15 | 341011 | BD Biosciences |
| Purity check | mouse anti-human CD127 | AF647 | HIL-7R-M21 | 0.625 | 560905 | BD Biosciences |
| Purity check | mouse anti-human CD4 | BV510 | SK3 | 1.25 | 562970 | BD Biosciences |

**Table S2:** Basal characteristics of patients with chronic coronary syndrome and healthy controls

|  | | |  | **Patients with CCS (n=20)** | | **Healthy controls (n=19)** | ***p*-value** |
| --- | --- | --- | --- | --- | --- | --- | --- |
| Age, years | | |  | | 65 (60 – 69) | 64 (60 – 70) | 0.756 |
| Female, n(%) |  | | | | 10 (50) | 10 (53) | 0.869 |
| Body mass index, kg*m^-2^ | |  | | | 28.7 (26.8 – 31.7) | 28.3 (26.6 – 30.4) | 0.877 |
| Smoking |  | | | |  |  |  |
| Current smoker, n(%) |  | | | | 2 (10) | 0 (0) | 0.157 |
| Former smoker, n(%) |  | | | | 6 (30) | 7 (37) | 0.651 |
| Diabetes type 2, n(%) |  | | | | 5 (25) | 0 (0) | 0.020 |
| Coronary artery disease severity (1-/2-/3-vessel disease), n(%) |  | | | | 6 (30)/7 (35)/7 (35) | - | - |
| Prior coronary intervention, PCI/CABG, n(%) |  | | | | 18 (90)/2 (10) | - | - |
| Left ventricular function, normal/mildly reduced/reduced^1^, n(%) |  | | | | 14 (70)/3 (15)/3 (15) | - | - |
| **Medication** |  | | | |  |  |  |
| Beta-blockers, n(%) |  | | | | 15 (75) | 0 (0) | <0.001 |
| Calcium channel blockers, n(%) |  | | | | 8 (40) | 0 (0) | 0.002 |
| ACEI/ARB, n(%) |  | | | | 18 (90) | 5 (26) | <0.001 |
| Antidiabetic drugs, n(%) |  | | | | 5 (25) | 0 (0) | 0.020 |
| Statins, n(%) |  | | | | 20 (100) | 2 (11) | <0.001 |
| Low-dose aspirin, n(%) |  | | | | 20 (100) | 0 (0) | <0.001 |
| **Laboratory measurements** |  | | | | | | |
| LDL cholesterol, mmol/L |  | | | | 1.40 (1.00 – 1.70) | n/a |  |
| HDL cholesterol, mmol/L |  | | | | 1.30 (1.20 – 1.60) | n/a |  |
| Triglycerides, mmol/L |  | | | | 1.10 (0.68 – 1.40) | n/a |  |
| Fasting glucose, mmol/L |  | | | | 5.4 (5.2 – 6.0) | n/a |  |
| CRP, mg/l |  | | | | 0.95 (0.43 – 1.70) | 1.10 (0.80 – 2.90) | 0.151 |
| IL-6, pg/ml |  | | | | 3.34 (2.21 – 4.94) | 2.58 (1.68 – 3.92) | 0.109 |
| TNF, pg/ml |  | | | | 17.9 (15.2 – 21.0) | 15.2 (12.0 – 17.1) | 0.011 |
| IFN-γ, pg/ml |  | | | | 0.25 (0.16 – 0.54) | 0.22 (0.17 – 0.37) | 0.518 |
| IL-18, pg/ml |  | | | | 279 (189 – 418) | 237 (197 – 314) | 0.448 |
| IL-1β, pg/ml |  | | | | 0.08 (0.06 – 0.16) | 0.11 (0.07 – 0.17) | 0.431 |
| IL-10, pg/ml |  | | | | 6.75 (5.84 – 8.71) | 5.26 (4.58 – 7.02) | 0.003 |
| MMP-8, pg/ml |  | | | | 867 (579 – 1139) | 677 (537 – 1113) | 0.457 |
| MMP-9, ng/ml |  | | | | 24.8 (19.2 – 40.4) | 36.4 (27.5 – 39.4) | 0.114 |
| MPO, ng/ml |  | | | | 49.1 (39.9 – 64.5) | 44.2 (34.8 – 63.8) | 0.238 |
| Total leukocyte count, 10^6^/mL |  | | | | 5.5 (4.9 – 6.9) | 5.4 (4.8 – 6.5) | 0.439 |
| Neutrophil count, 10^6^/mL |  | | | | 2.9 (2.4 – 4.0) | 3.3 (2.1 – 3.5) | 0.593 |

Values are given as n (%), or median (interquartile range), *p*-values are derived from Pearson Chi-Square Test or Mann-Whitney U Test, respectively. CCS, chronic coronary syndrome; PCI, percutaneous coronary intervention; CABG, coronary artery by-pass grafting; ACEI, angiotensin converting enzyme inhibitors; ARB, angiotensin receptor blockers; CRP, C-reactive protein; LDL, low density lipoprotein; n/a, not available; HDL, high density lipoprotein; IL, interleukin; TNF, tumor necrosis factor; IFN, interferon; MMP, matrix metalloprotease; MPO, myeloperoxidase. ^1^echocardiographic measurement of left ventricular ejection fraction, normal ≥ 50%, mildly reduced 41-49%, reduced ≤ 40%.

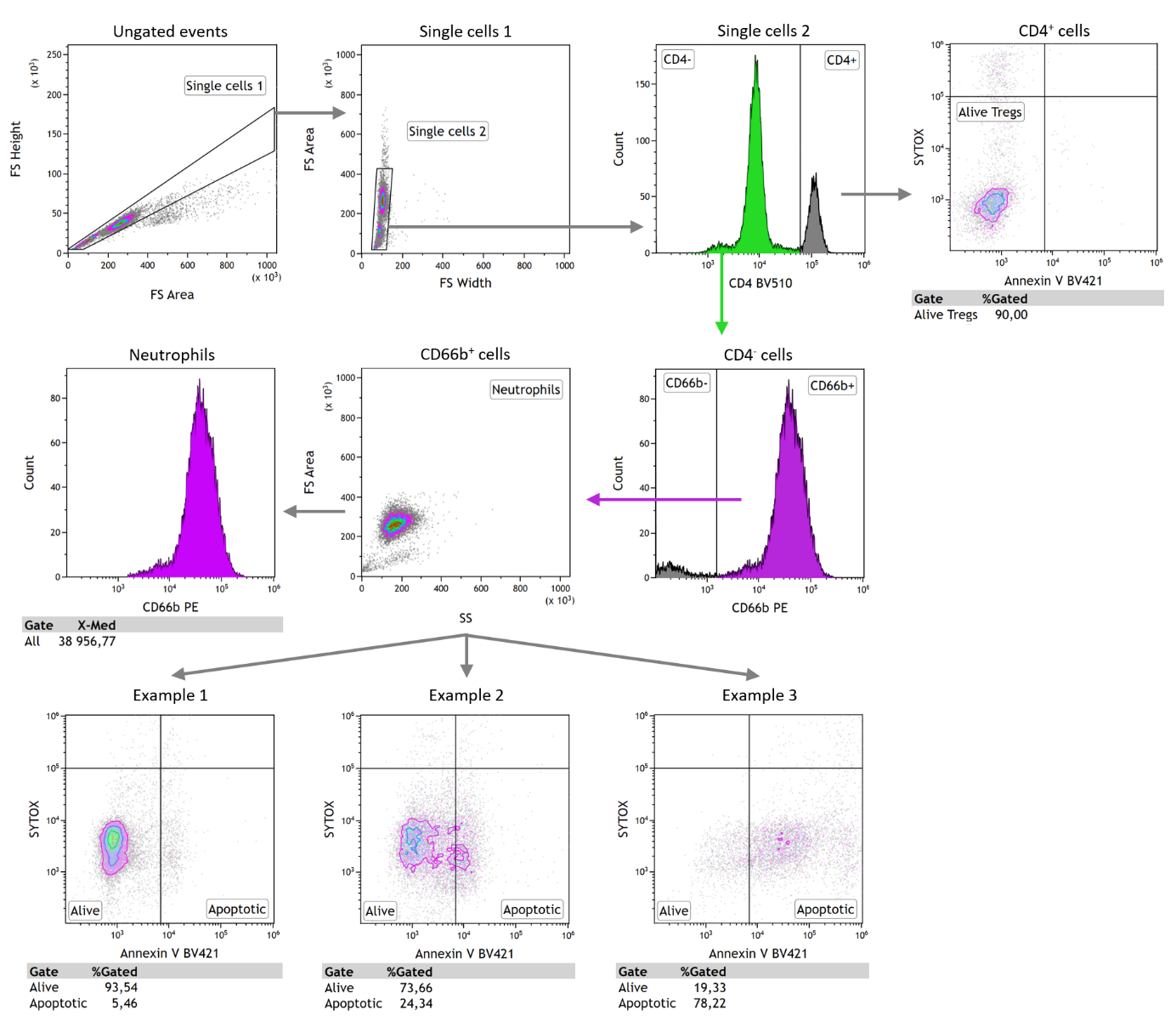

**Figure S1: Gating strategy to assess activation and apoptosis of neutrophils.**

Neutrophils were selected as CD4^-^, CD66b^+^ single cells. Activation was assessed by median value of CD66b mean fluorescent intensity (MFI). Apoptosis was assessed by determining the percentage of Annexin V^+^/SYTOX^-^ cells. Representative examples of one patients with chronic coronary syndrome (CCS) and two healthy controls are shown. T_reg_/CD4^+^ T cell survival was assessed by determining the percentage of Annexin V^-^/SYTOX^-^ CD4^+^ single cells.

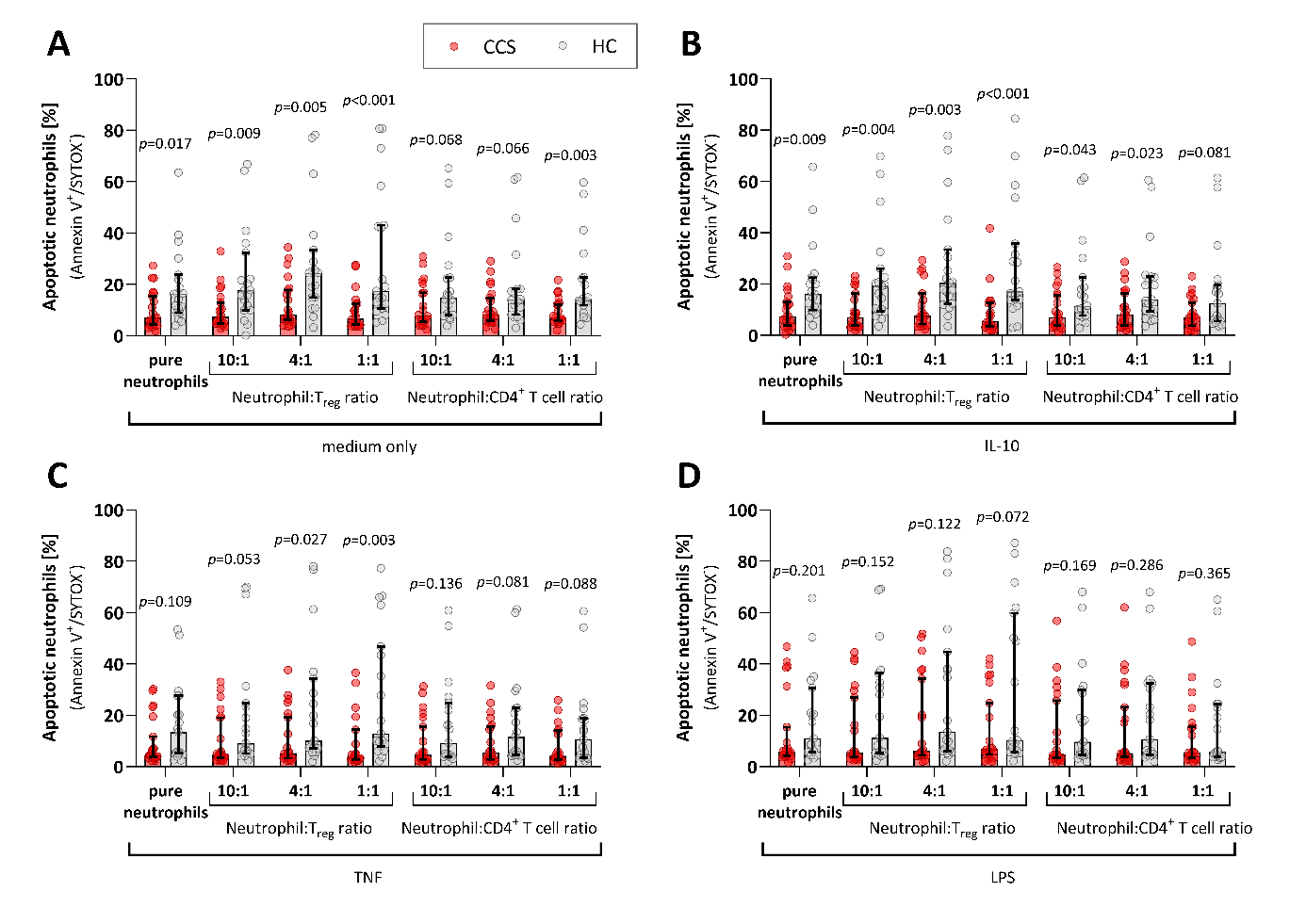

**Figure S2:** **T_regs_ induced neutrophil apoptosis in** **healthy controls, but not in patients with chronic coronary syndrome.**

T_regs_, CD4^+^ T cells and neutrophils were isolated from 20 patients with chronic coronary syndrome (CCS) and 19 healthy controls (HC). Neutrophil apoptosis was assessed by flow cytometry (% of Annexin V^+^/SYTOX^-^ cells) after 5 h co-culture with T_regs_ or CD4^+^ T cells at different ratios (neutrophils: T_regs_/CD4^+^ T cells 1:0, 10:1, 4:1 and 1:1) in medium only (**A**) or medium with interleukin (IL)-10 (20 ng/ml) (**B**), tumor necrosis factor (TNF, 1 ng/ml) (**C**) or lipopolysaccharide (LPS, 100 ng/ml) (**D**). Significances are between CCS and HC and values are given as median ± 95% confidence interval.

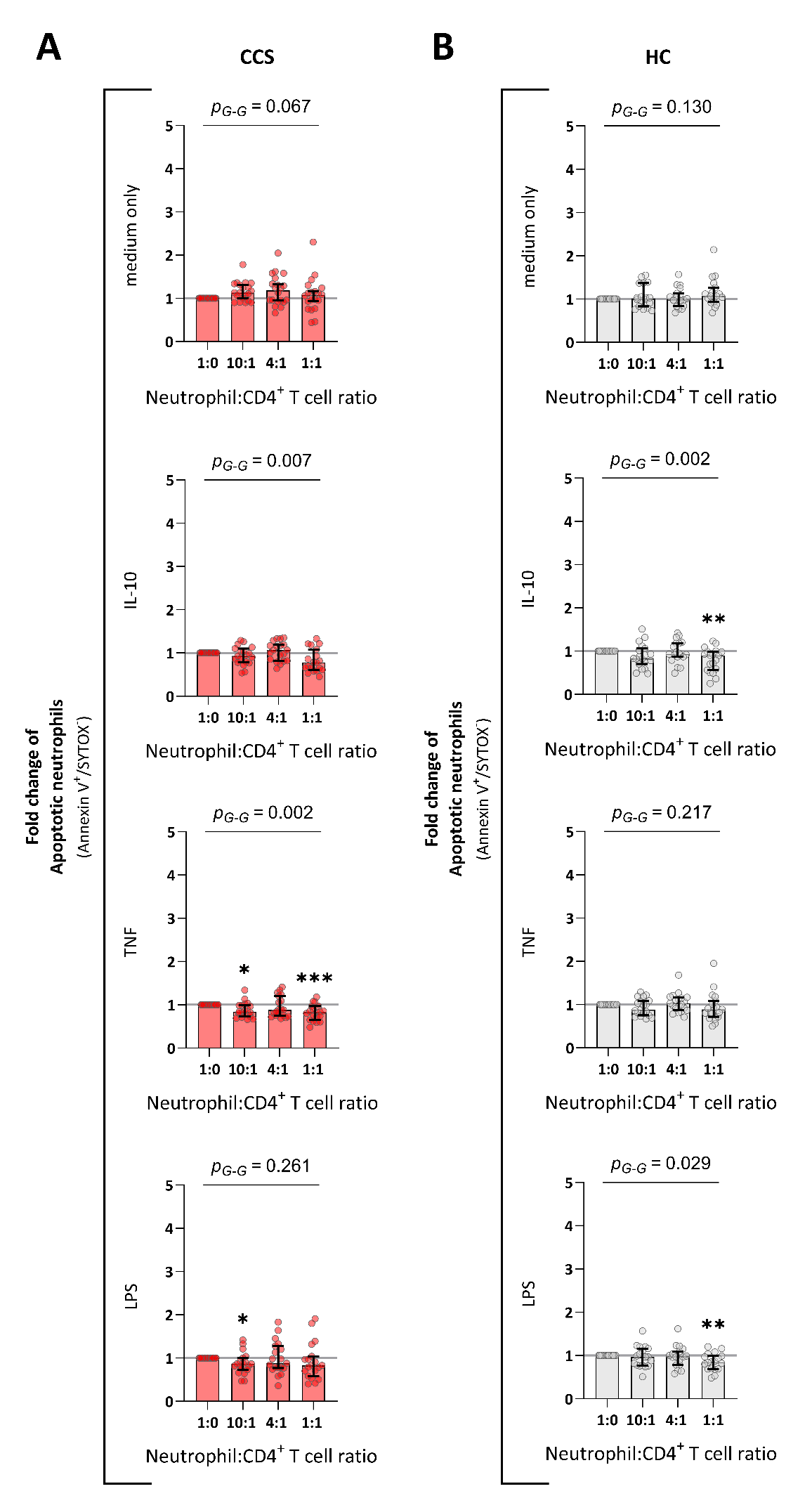

**Figure S3:** **The effect of neutrophil-CD4^+^ T cell co-cultures on neutrophil apoptosis in patients with chronic coronary syndrome and healthy controls.**

Neutrophils and CD4^+^ T cells were isolated from 20 patients with chronic coronary syndrome (CCS) and 19 healthy controls (HC). The cells were co-cultured at different ratios (neutrophils: CD4^+^ T cells 1:0, 10:1, 4:1 and 1:1) in medium only or medium with interleukin (IL)-10 (20 ng/ml), tumor necrosis factor (TNF, 1 ng/ml) or lipopolysaccharide (LPS, 100 ng/ml). After 5 h of co-culture, neutrophil apoptosis (Annexin^+^/SYTOX^-^) was determined by flow cytometry. The percentages of apoptotic neutrophils are presented as fold change compared to pure neutrophil culture in patients with CCS (**A**) and healthy controls (HC) (**B**). Significances are compared to 1:0 culture within CCS and HC groups, respectively, *p*-values are adjusted for multiple comparison via Dunn; CCS (**A**): **p*=0.030 in TNF ratio 10:1, ****p*<0.001 in TNF ratio 1:1, **p*=0.017 in LPS ratio 10:1; HC (**B**): ***p*=0.001 in IL-10 ratio 1:1, ***p*=0.006 in LPS ratio 1:1; and significance for trend is given as *p*-value of a Greenhouse-Geisser test. Values are given as median ± 95& confidence interval.

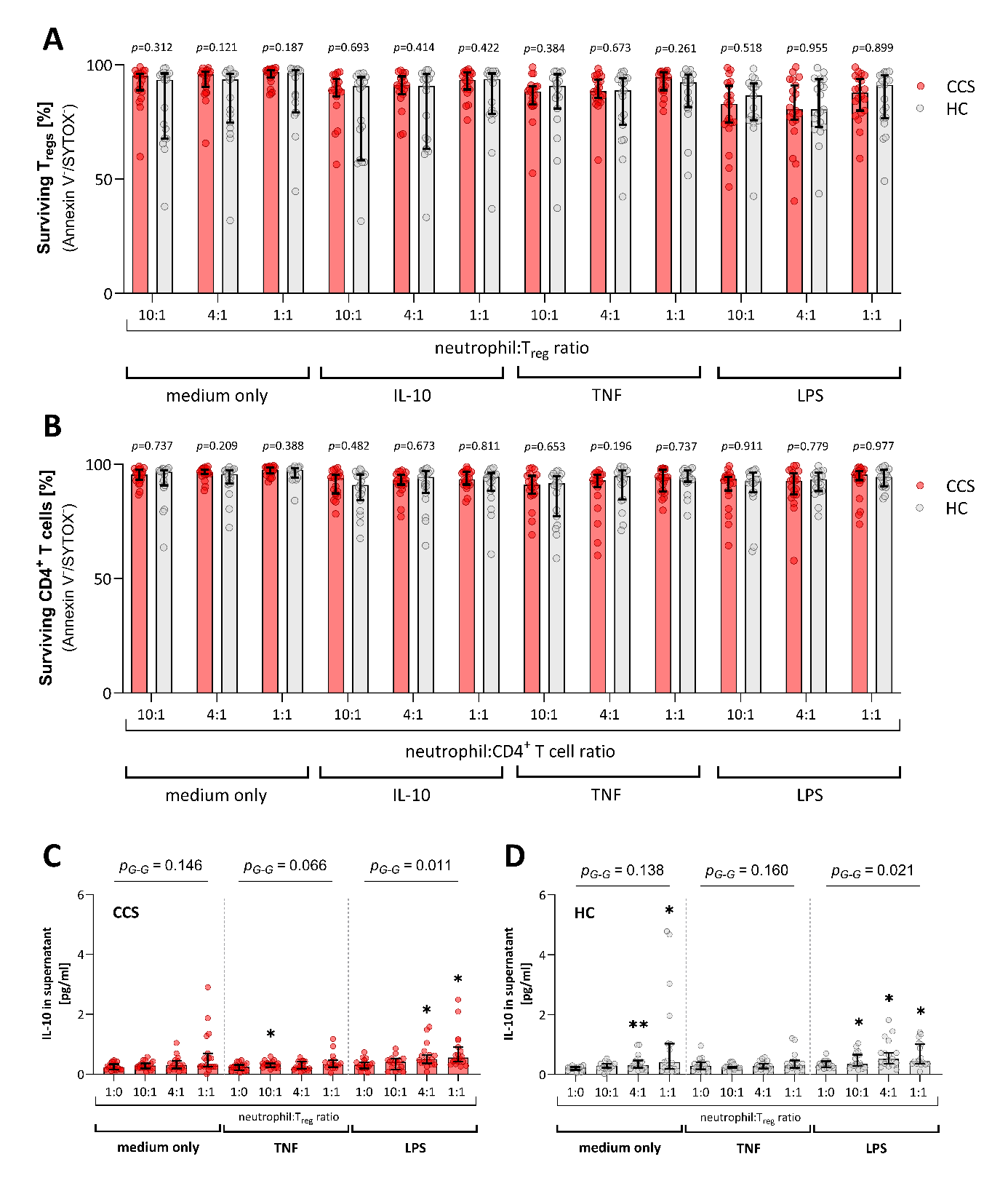

**Figure S4: T_reg_ survival and IL-10 release in patients with chronic coronary syndrome and healthy controls.**

The survival of T_regs_ and CD4+ T cells in neutrophil co-cultures was assessed by flow cytometry with Annexin V^-^/SYTOX^-^ cells deemed as surviving in samples from 20 patients with chronic coronary symptom (CCS) and 19 healthy controls (HC). The percentages of surviving T_regs_ (**A**) and CD4+ T cells (**B**) after 5 h of co-culture in medium only or in medium with interleukin (IL)-10 (20 ng/ml), tumor necrosis factor (TNF, 1 ng/ml) or lipopolysaccharides (LPS, 100 ng/ml) are shown. Significances are between CCS and HC and values are presented as median ± 95% confidence interval.

Concentrations of IL-10 were measured by Luminex in supernatants from neutrophil-T_reg_-co-cultures after 5h incubation in medium only or in medium with TNF (1 ng/ml) or LPS (100 ng/ml) in 20 patients with CCS (**C**) and in HC (**D**). Significances are between pure neutrophils and neutrophil-T_reg_-cocultures within CCS and HC groups, respectively; p-values were adjusted for multiple comparison via Dunnett; CCS (**C**): **p*=0.049 in TNF ratio 10:1, **p*=0.023 in LPS ratio 4:1, **p*=0.019 in LPS ratio 1:1; HC (**D**): ***p*=0.006 in medium only ratio 4:1, **p*=0.020 in medium only ratio 1:1, **p*=0.042 in LPS ratio 10:1, **p*=0.012 in LPS ratio 4:1, **p*=0.022 in LPS ratio 1:1; and significance for trend is given as *p*-value of a Greenhouse-Geisser test. Values are presented as median ± 95% confidence interval.
